# The Pupillary Light Response Reflects Visual Working Memory Content

**DOI:** 10.1101/477562

**Authors:** Cecília Hustá, Edwin Dalmaijer, Artem Belopolsky, Sebastiaan Mathôt

**Affiliations:** Faculty of Science and Engineering, University of Groningen, The Netherlands; Department of Experimental Psychology, University of Groningen, The Netherlands; MRC Cognition and Brain Sciences Unit, University of Cambridge, United Kingdom; Department of Experimental and Applied Psychology, VU University Amsterdam, Netherlands

**Keywords:** pupillometry, pupil light response, visual working memory

## Abstract

Recent studies have shown that the pupillary light response (PLR) is modulated by higher cognitive functions, presumably through activity in visual sensory brain areas. Here we use the PLR to test the involvement of sensory areas in visual working memory (VWM). In two experiments, participants memorized either bright or dark stimuli. We found that pupils were smaller when a pre-stimulus cue indicated that a bright stimulus should be memorized; this reflects a covert shift of attention during encoding of items into VWM. Crucially, we obtained the same result with a post-stimulus cue, which shows that internal shifts of attention within VWM affect pupil size as well. Strikingly, pupil size reflected VWM content only briefly. This suggests that a shift of attention within VWM momentarily activates an “active” memory representation, but that this representation quickly transforms into a “hidden” state that does not rely on sensory areas.

## The Pupillary Light Response Reflects Visual Working Memory Content

Traditionally, the pupillary light response (PLR) was considered a reflex in response to changes in environmental brightness. However, recent studies have demonstrated that the PLR is modulated by higher-level cognition (reviewed in Binda & Murray, 2015; Mathôt, 2018). Such effects likely occur when higher-level cognition affects activity in visual sensory brain areas, which is subsequently “read out” by the pupils.

For example, in several studies, participants were presented with both dark and bright stimuli. Participants were subsequently cued to attend to either the bright or the dark stimulus, without shifting their gaze (i.e. covert attention). Attending to the bright stimuli resulted in smaller pupils than attending to the dark stimuli (Binda, Pereverzeva, & Murray, 2013; Mathôt, van der Linden, Grainger, & Vitu, 2013; Naber, Alvarez, & Nakayama, 2013). Single-cell-recording studies have linked this effect to the frontal eye fields (FEF), a part of the frontal cortex that is associated with covert attention. Microstimulation of FEF results in increased attention to a specific part of the visual field (Moore & Fallah, 2001). Crucially, if the stimulated region corresponds to the location where a bright stimulus appears, the pupil constricts more strongly than if the stimulus appears at a different, unstimulated location (Ebitz & Moore, 2017). A similar effect has been reported for microstimulation of the superior colliculus (SC), a midbrain region that is also associated with visual attention (Wang & Munoz, in press). Taken together, both behavioral and neurophysiological studies have shown that covert visual attention enhances the PLR.

A PLR can even be elicited without the physical presence of bright or dark stimuli. In studies of mental imagery, participants were instructed to imagine stimuli that had previously been presented with varying brightness levels. The size of the pupil varied depending on the imagined brightness, with brighter objects resulting in smaller pupils (Laeng & Sulutvedt, 2014). This effect was replicated with mental imagery of real-life scenarios: Imagery of scenes like “a sunny sky” resulted in smaller pupils than imagery of scenes like “a dark room” (Laeng & Sulutvedt, 2014). These results are consistent with the finding that similar visual sensory areas are active during perception and mental imagery of visual objects (Ganis, Thompson, & Kosslyn, 2004). Presumably, the activity in visual sensory areas that is elicited by mental imagery subsequently affects pupil size.

According to many theoretical frameworks, mental imagery is highly related to visual working memory (VWM). VWM is a system with limited storage capacity that holds visual information ready for immediate use. VWM consists of encoding and maintenance. During encoding, visual stimuli are visible and a VWM representation is created (Bundesen, 1990; Dalmaijer, Manohar, & Husain, 2018). During maintenance, stimuli are no longer visible, and their VWM representations therefore need to be rehearsed so that they can be used later (Zokaei, Heider, & Husain, 2014). Analogous to mental imagery, maintaining of stimuli in VWM activates visual sensory areas (Yi, Turk-Browne, Chun, & Johnson, 2008). The similarity between mental imagery and VWM maintenance leads to the prediction that maintaining bright stimuli in VWM should lead to pupil constriction.

However, the only study so far that investigated this question reported that maintaining bright or dark stimuli in VWM did *not* affect the PLR (Blom, Mathôt, Olivers, & Van der Stigchel, 2016). Blom and colleagues (2016) cued participants to memorize either bright or dark objects. In different experiments, participants memorized the shape, orientation, or the exact brightness level of the stimuli. Pupil size was significantly smaller when participants were encoding the bright as compared to the dark stimuli. However, this effect faded approximately one second after the stimuli disappeared from the screen. This led Blom and colleagues (2016) to conclude that the PLR reflects VWM content during encoding, but not during maintenance. Phrased differently, the authors concluded that keeping bright or dark objects in VWM does not affect pupil size.

However, there are several alternative explanations for the results of Blom and colleagues (2016) that warrant a re-investigation of the question. Notably, in their experiments, participants were presented with a cue *before* (rather than after) the presentation of the brightness-related stimuli, and this cue indicated whether only the bright or only the dark stimuli needed to be memorized. Therefore, participants covertly shifted their attention to either the bright or the dark stimuli while these were actually present on the screen, leading to differences in pupil size (cf. Binda et al., 2013; Mathôt et al., 2013). Crucially, because this pupil-size difference persisted into the maintenance period, it was not clear whether any pupil-size differences during maintenance were due to VWM per se, or merely reflected a carry-over effect from the encoding phase (as Blom and colleagues concluded).

The question of whether brightness-related content in VWM affects pupil size is important, because, if so, this would strongly suggest that VWM relies on sensory brain areas—currently a hotly debated topic (Gayet, Paffen, & Van der Stigchel, 2018; Xu, 2017). Therefore, to firmly establish whether keeping bright or dark stimuli in VWM affects pupil size, we designed a study that allowed us to distinguish any effects due to VWM encoding from effects due to VWM maintenance. In one condition of our experiment, we wanted to replicate the effect of covert visual attention on the PLR (Binda et al., 2013; Blom et al., 2016; Mathôt et al., 2013). As shown in many earlier studies, we expected that directing covert attention to dark or bright stimuli during VWM encoding would be reflected in the PLR. In another condition, we introduced a retro-cue to investigate the relationship between VWM maintenance and the PLR (cf. Belopolsky & Theeuwes, 2011). Participants were instructed to first encode both bright and dark stimuli, so that the encoding phase was identical in all trials. Subsequently, participants dropped one stimulus from VWM when the retro-cue was presented, leaving either only bright or only dark stimuli for maintenance in VWM. Crucially, we predicted that maintaining bright stimuli would result in smaller pupils as compared to maintaining dark stimuli.

## Results

### Experiment 1

Experiment 1 included two conditions: Pre-Cue and Retro-Cue (Figure 1a & 1c). In the Pre-Cue condition, participants were cued to either attend to the stimulus appearing on the left or right side. Next, they were presented with one bright and one dark circle. The participants’ task was to maintain the precise brightness level of the cued circle. Finally, after a retention interval of 4 s, participants indicated whether the brightness of a newly presented circle was the same as, or different from, the brightness of the memorized circle. The Retro-Cue Condition was almost identical to the Pre-Cue Condition, with the important difference that the to-be-memorized stimuli were shown before the cue. We used a Quest adaptive procedure (Watson & Pelli, 1983) to ensure that the response accuracy for both bright and dark stimuli was kept at around 75%. In the Pre-Cue Condition, the mean accuracy was 75% for bright trials and 74% for dark trials. In the Retro-Cue Condition, the mean accuracy was 71% for bright and 69% for dark trials.

**Figure 1.**
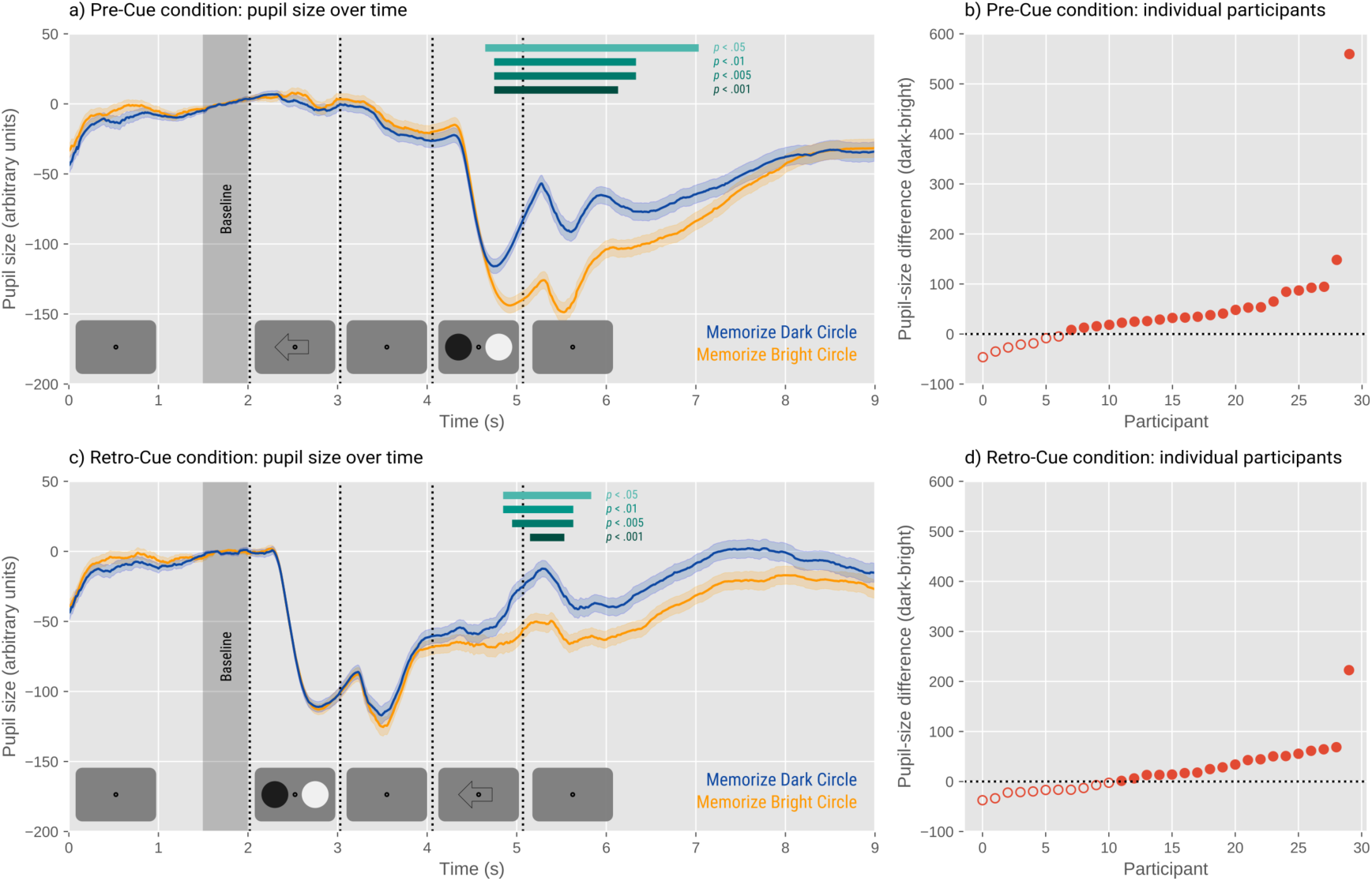
Results of Experiment 1. a) This figure shows averaged pupil size for all participants through the progression of all Pre-Cue trials. The orange line represents the average pupil size when bright stimuli are the targets and the blue line when dark stimuli are the targets. The shaded error bands represent the grand standard error (i.e. across individual trials). b) Shows the average effects of individual participants in the Pre-Cue Condition between 4000 ms and 6000 ms calculated by subtracting the mean pupil size for bright trials from dark trials. c) Shows averaged pupil size for all participants through the progression of all Retro-Cue trials. d) Shows the average effects of individual participants in the Retro-Cue Condition calculated in the same way as for the Pre-Cue Condition.

We conducted a linear mixed effects analysis (LME) on all trials (correct and incorrect; analyzing only correct trials did not qualitatively change the results) with Pupil Size as dependent measure, and two fixed effects, each containing two levels (Brightness: Bright and Dark; Condition: Pre-Cue and Retro-Cue). We include by-participant random intercepts and slopes for all fixed effects. This analysis was conducted for each 10 ms time window separately. We considered effects significant if *t* > 1.96 (cf. Mathot et al., 2014; Mathôt et al., 2013), although we emphasize overall patterns and effect sizes rather than significance of individual data points.

There was a significant interaction between Brightness and Condition between 5100 ms and 5590 ms, indicating that right at the start of the maintenance phase, the effect of brightness depended on the condition (Pre-Cue or Retro-Cue). Subsequently, we performed two separate LMEs for the two conditions (also run with all trials).

For the Pre-Cue Condition, there was a significant effect of Brightness from 4700 ms to 6990 ms, meaning that the pupil difference appeared during encoding of the brightness-related stimuli and briefly persisted into the maintenance phase (Figure 1a). Twenty-three participants (out of 30) exhibited the effect in the expected direction (Figure 1b). In general, this means that when participants covertly attended to the white circles on the encode screen, their pupils were smaller than when they attended to the black circles.

In the Retro-Cue Condition, there was an effect of Brightness from 4900 ms until 5790 ms (Figure 1c), directly corresponding to the maintenance phase. The effect occurred in the expected direction for 19 participants (out of 30; Figure 1d). This indicates that VWM content is reflected in the PLR not only during encoding but also during maintenance. Phrased differently, shifting attention within VWM representations (that are brightness-related) is reflected in pupil size, such that internally shifting attention toward bright stimuli elicits smaller pupils than internally shifting attention toward dark stimuli.

In summary, by using a retro-cue to shift attention within vWM representations, we were able to examine the relationship between maintenance of brightness-related content and pupil size while keeping stimulus encoding constant. Our results clearly show that shifting attention to bright stimuli in vWM results in pupil constriction relative to maintaining dark stimuli.

### Experiment 2

In Experiment 2, we investigated if the relationship between the VWM content and the PLR depends on whether VWM representations are in a high- or low-priority state, following single-item-template theories that postulate that only a single VWM item can be in a high-priority state at a time, and that only this item is represented in visual sensory areas (Folk & Anderson, 2010; Houtkamp & Roelfsema, 2009; Oberauer, 2002; Olivers, Peters, Houtkamp, & Roelfsema, 2011; Zokaei, Manohar, Husain, & Feredoes, 2014), and thus affects the size of the pupil. We investigated this by varying memory load. Participants were either asked to maintain one item or two items during the retention interval. Single-item-template theories would predict that the PLR would reflect VWM content only when participants maintained one item, which was in a high-priority state, and not when participants maintained two items, which would then compete with each other and both take on a low-priority state (cf. Olivers et al., 2011; van Moorselaar, Theeuwes, & Olivers, 2014).

We designed two conditions: Set-Size-One and Set-Size-Two. The Set-Size-One Condition was an exact replication of the Retro-Cue-Condition (Figure 2a). In the Set-Size-Two Condition, we presented participants with four stimuli during the encoding phase. The retro cue indicated whether the two circles on the right or on the left were task relevant. During the response phase, participants had to indicate whether the two new circles were both identical to the memorized circles, or if one of them was different (Figure 2c). A Quest adaptive procedure (Watson & Pelli, 1983) controlled for accuracy separately for the Bright and Dark Conditions as well as for the Set-Size-One and Set-Size-Two Conditions to try to keep accuracy at a constant 75% in all conditions. In the Set-Size-One Condition, the mean accuracy for the bright trials was 69% and 68% for the dark trials. In the Set-Size-Two Condition, the accuracy was 74% for the bright and 71% for the dark trials.

**Figure 2.**
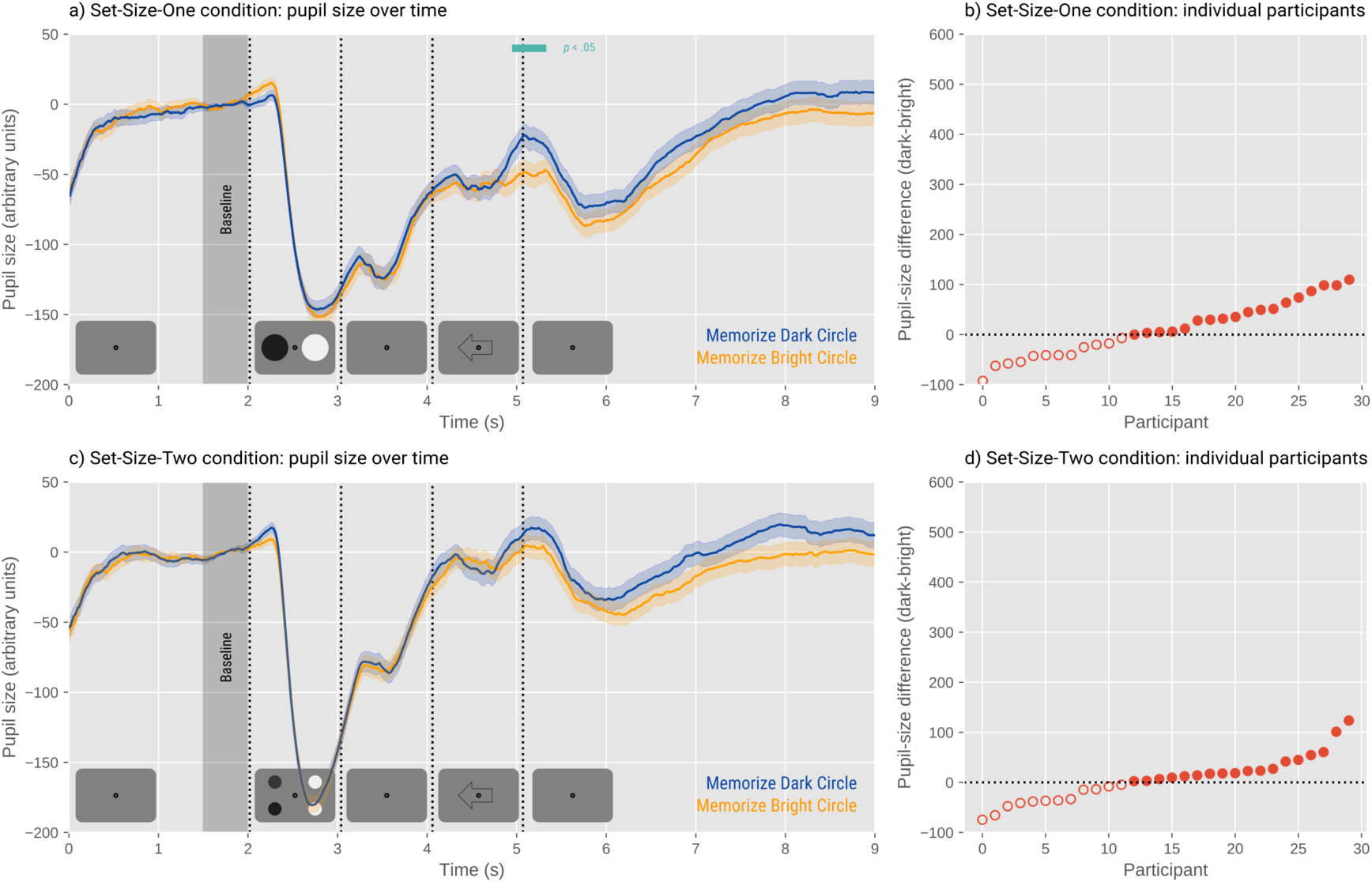
Results of Experiment 2. a) This figure shows averaged pupil size for all participants through the progression of all Set-Size-One trials. The orange line represents the average pupil size when bright stimuli are the targets and the blue line when dark stimuli are the targets. The shaded error bars represent the grand standard error (i.e. across individual trials). b) Shows the average effects of individual participants in the Set-Size-One Condition between 4000 ms and 6000 ms calculated by subtracting the summed pupil size for bright trials from dark trials. c) Shows averaged pupil size for all participants through the progression of all Set-Size-Two trials. d) Shows the average effects of individual participants in the Set-Size-Two Condition calculated in the same way as for the Pre-Cue Condition.

A similar analysis was performed as for Experiment 1, using Brightness (Bright and Dark) and Memory Load (Set-Size-One and Set-Size-Two) as fixed effects. This analysis revealed no interaction between Memory Load and Brightness after the presentation of the retro cue. This means that the effect of brightness on the PLR did not notably differ between the two memory-load conditions. This suggests that the effect of brightness-related VWM content on the PLR does not crucially depend on the priority status of the items in VWM. (There was a spurious interaction between Brightness and Condition from 2100 ms to 2290 ms, which corresponded to the encoding phase. However, no interaction could occur before presentation of the targets, as participants were not aware which brightness would have to be encoded, and hence this effect is necessarily spurious.)

Despite not finding a significant interaction in the overall model, we analyzed the two Memory Load conditions separately. In the Set-Size-One Condition, we replicated the significant effect of Brightness on pupil size between 5000 ms and 5290 ms, meaning that on average, the participants’ pupils were smaller when maintaining bright circles as compared to the dark at the beginning of the maintenance phase (Figure 3a). However, it should be noted that this effect was weaker and present for fewer (*N* = 18, of 30) participants than in Experiment 1.

The Set-Size-Two Condition revealed no significant effects of brightness on pupil size (Figure 2c). However, because the effect was qualitatively in the same direction as for the Set-Size-One Condition, and because the interaction between Brightness and Memory Load was not reliable, we do not draw any conclusions about an effect of Memory Load. Eighteen participants (out of 30) had smaller pupils when maintaining bright stimuli as compared to the dark in the Set-Size-Two Condition (Figure 2d).

## Discussion

In two experiments, we examined whether visual working memory (VWM) content is reflected in the pupillary light response (PLR). Specifically, we wanted to know whether maintaining bright stimuli in VWM is associated with smaller pupils than maintaining dark stimuli. Overall, we showed that VWM content is reflected in the PLR during both encoding and maintenance. Consistent with previous studies (Binda et al., 2013; Laeng & Sulutvedt, 2014; Mathôt et al., 2013), this shows that the PLR, which was previously thought of as a simple reflex, is controlled by higher cognitive processes, such as working memory. Our results further suggest that VWM involves sensory representations, presumably in visual cortex (Yi et al., 2008), which subsequently trigger pupil responses.

A striking aspect of our results is the time course (in the Retro-Cue Condition of Experiment 1, and the Set-Size-One Condition of Experiment 2). The content of VWM affected pupil size only briefly after the presentation of the retro-cue, rather than throughout the entire retention interval. This was unexpected, considering that we anticipated that this effect reflected maintenance of different brightness levels, which should occur during the entire retention interval. However, this finding is consistent with recent studies showing that VWM maintenance is not accompanied by sustained activity in visual sensory brain areas, but rather that such activity is periodical or transient (Rose et al., 2016; Sprague, Ester, & Serences, 2016; Sreenivasan, Curtis, & D’Esposito, 2014; Stokes, 2015; Wolff, Jochim, Akyürek, & Stokes, 2017). Our finding that pupil size reflects VWM content only briefly may reflect a transition from an ‘active’ state (which is reflected in pupil size) to a ‘hidden’ state (which is not reflected in pupil size). This provides unique new support for the notion of hidden VWM states, which has so far come primarily from decoding analyses in brain imaging; however, decoding studies provide inconclusive evidence for hidden states, because simulations indicate that re-emerging stimulus decodability in neuroimaging data could also reflect sustained neural activity (Schneegans & Bays, 2017).

We did not find a compelling dissociation between maintenance of one or two brightness-related stimuli. Such a dissociation would be predicted by strong single-item-template theories, which hold that there can be only one active item in VWM at a time, and that competition within VWM representations avoids any item from becoming active when multiple items are kept in VWM (van Moorselaar et al., 2014). We found qualitatively similar, but weaker effects on pupil size with a memory load of two items, as compared to one item. Overall, our results suggest that whether a VWM item is in an active or “silent” state depends strongly on time, and at most weakly on memory load.

So why is pupil size affected by the content of VWM and other cognitive factors? Possibly, the effect of higher cognitive functions on pupil size are preparatory mechanisms that optimize pupil size in anticipation of an environmental change in brightness (Mathôt, van der Linden, Grainger, & Vitu, 2015). For example, if you are in the dark and think about turning on a lamp, it is likely that you are going to do this soon. Therefore, it might be beneficial for your pupils to constrict before a sudden change of luminance impairs your vision.

## Methods

Participant data, experimental scripts, and analysis scripts are available from https://osf.io/ejxfa/.

### Experiment 1

#### Participants

We recruited 30 first-year psychology students from the University of Groningen, who participated in the present study for course credit. All participants had normal or corrected to normal vision, except for two participants, who took part in the experiment without their glasses, as calibration was not possible with them. The age of the participants ranged between 18 and 54 (*M* = 21, *SD* = 6.47), and 23 participants were females, six were males, and one identified as different gender. Both experiments were approved by the local ethics review board of the Department of Psychology of the University of Groningen (17370-S-NE).

#### Apparatus

Participants’ eye movements and pupil sizes were recorded with an Eyelink 1000 (SR Research, Mississauga, Canada, ON), and the data was sampled at 1000 Hz. The data was collected by recording the size of the right pupil of all of the participants. The collection was done in a dark room and participants were asked to place their head in a chin rest throughout the experiment. The task was designed with OpenSesame 3.2.0 (Mathôt, Schreij, & Theeuwes, 2012), using the PyGaze plug-ins for eye tracking (Dalmaijer, Mathôt, & Van der Stigchel, 2014). The stimuli were presented on a monitor with LCD display with 60 Hz refresh rate and resolution of 1920 x 1080.

#### Procedure and Stimuli

Before the experiment started, the eye tracker was calibrated with a five-point calibration procedure. Afterward, participants took part in a task in which they memorized a particular brightness level of black and white circles that appeared on a grey background (62 cd/m^2^). Participants were instructed to keep their eyes focused on a black fixation dot (2 cd/m^2^) that was in the center at all times. This was ensured by presenting a drift correction at the beginning of each trial, which paused the experiment unless participants shifted their gaze back to the center. Participants had a chance to get familiar with the task during a practice phase (10 trials). Experiment 1 was composed of 16 blocks, each lasting 3.47 minutes at most (excluding the duration of the drift corrections); the precise duration depended on the speed of responses. Each block consisted of 16 trials, with half the trials belonging to the Pre-Cue and half to the Retro-Cue Condition presented in random order.

In the Pre-Cue condition participants were initially presented with a cue (arrow pointing to the left or right) indicating whether the stimulus on the left or right would be task relevant. Subsequently two stimuli appeared (one black and one white circle), one of which they had to encode. This was followed by a retention interval lasting for 4 seconds. During the response phase, participants were presented with a circle of the same or a similar brightness as the one they had memorized. Participants had to report whether the brightness of this circle was the same as, or different from, the one they had memorized. The Retro-Cue Condition was almost identical to the Pre-Cue one, however, the order of the cue and target were reversed. (For durations of individual phases of the trial see Figure 1a and 1c.)

The targets for the bright and dark trials were selected from a specified brightness ranges. The bright range extended from 88 cd/m^2^ to 96 cd/m^2^, and the dark range extended from 11 cd/m^2^ to 19 cd/m^2^. A different response stimulus was brighter on some trials and darker on others. The size of this difference was controlled by a Quest adaptive procedure. It was implemented to control for participants’ accuracies, holding them constant at 75% for dark and bright stimuli separately.

After participants completed the task, they were asked about the strategies they used throughout the experiment (see supplementary materials at https://osf.io/ejxfa/).

#### Exclusion Criteria

For both conditions, trials in which the pupil at baseline was smaller than 2.1 mm in diameter or greater than 6.8 mm in diameter (*N*(trial) = 1) were excluded (as values above these were clear outliers based on a visual inspection of the pupil-baseline histogram). Additionally, in the Pre-Cue Condition, trials were excluded if participants horizontal gaze position deviated from the central band (between the targets) position during the presentation of the encode screen (*N*(trial) = 522). No such exclusion criteria were introduced for the Retro-Cue Condition, considering that participants did not know which stimulus was the target during the presentation of the encode screen, and eye movements could therefore not be systematically biased towards the to-be-memorizes stimulus.

### Experiment 2

#### Participants

Thirty participants were recruited from the same sample pool as in Experiment

1. All participants had normal or corrected to normal vision. The age of the participants ranged between 18 and 34 (*M* = 21.03, *SD* = 3.00), and 19 were females and 11 were males.

#### Apparatus

The same setup as in Experiment 1 was used.

#### Procedure and Stimuli

Identical calibration and drift correction procedures were used as in Experiment 1. The goal of the task was again to remember brightness level of black and white stimuli, but the number of stimuli varied. Participants had a chance to get familiar with the task during a practice phase (16 trials). Experiment 2 was composed of 14 blocks, each again lasting at most 3.47 minutes (excluding the duration of the drift corrections). Each block consisted of 16 trials; in eight trials participants had to maintain one stimulus (Set-Size-One Condition) and in the other eight trials they had to maintain two stimuli (Set-Size-Two Condition) in their VWM. The stimuli were dark on half of the trials and bright on the other half, all appearing on grey background (62 cd/m^2^). The conditions were presented in random order within blocks.

The sequence of both conditions was the same as in the Retro-Cue Condition in Experiment 1. The Retro-Cue Condition from Experiment 1 was identical to the Set-Size-One Condition in Experiment 2, in which participants had to maintain one stimulus in VWM (Figure 2a). In Set-Size-Two Condition, participants had to maintain two stimuli in their VWM after four circles were presented on the encode screen (Figure 2c). The subsequent arrow indicated whether the stimuli on the right or left were task relevant. Two circles were presented on the response screen. On the different trials the brightness of only one of the circles changed to ensure that participants were remembering the brightness of both circles.

The targets for the bright and dark trials were again selected from a specified brightness ranges. There were two bright ranges (very bright: 101 cd/m^2^ – 113 cd/m^2^, somewhat bright: 83 cd/m^2^ – 92 cd/m^2^) and two dark ranges (somewhat dark: 23 cd/m^2^ – 32 cd/m^2^, very dark: 2 cd/m^2^ – 13 cd/m^2^). How a stimulus was changed in different trials depended on the brightness range it was selected from (very bright and somewhat dark always changed to a darker stimulus, somewhat bright and very dark always changed to a lighter stimulus). When four circles were presented on the screen, they were all selected from different brightness ranges, to ensure sufficient variance in the brightness levels. The size of the brightness difference was controlled by a Quest adaptive procedure. It was implemented to control for the participants’ accuracies, holding them constant at 75%, and it controlled for the accuracy separately in the four conditions (Set-Size-One Bright, Set-Size-One Dark, Set-Size-Two Bright, and Set-Size-Two Dark).

After the participants completed the task they were again asked about the strategies they used throughout the experiment (see supplementary materials for more information at https://osf.io/ejxfa/).

### Exclusion Criteria

Trials on which the pupil size at baseline was lower than 2.1 mm or higher than 6.8 mm in diameter (*N* = 10) were again excluded.

## References

Belopolsky, A. V., & Theeuwes, J. (2011). Selection within visual memory representations activates the oculomotor system. Neuropsychologia, 49(6), 1605–1610. https://doi.org/10.1016/j.neuropsychologia.2010.12.045

Binda, P., & Murray, S. O. (2015). Keeping a large-pupilled eye on high-level visual processing. Trends in Cognitive Sciences, 19(1), 1–3. https://doi.org/10.1016/j.tics.2014.11.002

Binda, P., Pereverzeva, M., & Murray, S. O. (2013). Pupil constrictions to photographs of the sun. Journal of Vision, 13(6), 1–9. https://doi.org/10.1167/13.6.8

Blom, T., Mathôt, S., Olivers, C. N. L., & Van der Stigchel, S. (2016). The pupillary light response reflects encoding, but not maintenance, in visual working memory. Journal of Experimental Psychology: Human Perception and Performance, 42(11), 1716–1723. https://doi.org/10.1037/xhp0000252

Bundesen, C. (1990). A theory of visual attention. Psychological Review, 97(4), 523–547. Retrieved from http://www.ncbi.nlm.nih.gov/pubmed/2247540

Dalmaijer, E. S., Manohar, S. G., & Husain, M. (2018). Parallel encoding of information into visual short-term memory. https://doi.org/398990.

Dalmaijer, E. S., Mathôt, S., & Van der Stigchel, S. (2014). PyGaze: An open-source, cross-platform toolbox for minimal-effort programming of eyetracking experiments. Behavior Research Methods, 46(4), 913–921. https://doi.org/10.3758/s13428-013-0422-2

Ebitz, B. R., & Moore, T. (2017). Selective modulation of the pupil light reflex by microstimulation of prefrontal cortex. The Journal of Neuroscience, 37(19), 5008–5018. https://doi.org/10.1523/JNEUROSCI.2433-16.2017

Folk, C. L., & Anderson, B. A. (2010). Target-uncertainty effects in attentional capture: Color-singleton set or multiple attentional control settings? Psychonomic Bulletin and Review, 17(3), 421–426. https://doi.org/doi:10.3758/PBR.17.3.421

Ganis, G., Thompson, W. L., & Kosslyn, S. M. (2004). Brain areas underlying visual mental imagery and visual perception: An fMRI study. Cognitive Brain Research, 20(2), 226–241. https://doi.org/10.1016/j.cogbrainres.2004.02.012

Gayet, S., Paffen, C. L. E., & Van der Stigchel, S. (2018). Visual working memory storage recruits sensory processing areas. Trends in Cognitive Sciences, 22(3), 189–190. https://doi.org/10.1016/j.tics.2017.09.011

Houtkamp, R., & Roelfsema, P. R. (2009). Matching of visual input to only one item at any one time. Psychological Research, 73(3), 317–326. https://doi.org/10.1007/s00426-008-0157-3

Laeng, B., & Sulutvedt, U. (2014). The eye pupil adjusts to imaginary light. Psychological Science, 25(1), 188–197. https://doi.org/10.1177/0956797613503556

Mathôt, S. (2018). Pupillometry: Psychology, physiology, and function. Journal of Cognition, 1(16), 1–23. https://doi.org/10.5334/joc.18

Mathot, S., Dalmaijer, E., Grainger, J., & Van der Stigchel, S. (2014). The pupillary light response reflects exogenous attention and inhibition of return. Journal of Vision, 14(14), 7–7. https://doi.org/10.1167/14.14.7

Mathôt, S., Schreij, D., & Theeuwes, J. (2012). OpenSesame: An open-source, graphical experiment builder for the social sciences. Behavior Research Methods, 44(2), 314–324. https://doi.org/10.3758/s13428-011-0168-7

Mathôt, S., van der Linden, L., Grainger, J., & Vitu, F. (2013). The pupillary light response reveals the focus of covert visual attention. PLoS ONE, 8(10). https://doi.org/10.1371/journal.pone.0078168

Mathôt, S., van der Linden, L., Grainger, J., & Vitu, F. (2015). The pupillary light response reflects eye-movement preparation. Journal of Experimental Psychology: Human Perception and Performance, 41(1), 28–35. https://doi.org/10.1037/a0038653

Moore, T., & Fallah, M. (2001). Control of eye movements and spatial attention. PNAS, 98(3), 1273–1276. https://doi.org/10.1073/pnas.98.3.1273

Naber, M., Alvarez, G. A., & Nakayama, K. (2013). Tracking the allocation of attention using human pupillary oscillations. Frontiers in Psychology, 4(3), 592–600. https://doi.org/10.3389/fpsyg.2013.00919

Oberauer, K. (2002). Access to information in working memory: Exploring the focus of attention. Journal of Experimental Psychology. Learning, Memory, and Cognition, 28(3), 411–421. Retrieved from http://www.ncbi.nlm.nih.gov/pubmed/12018494

Olivers, C. N. L., Peters, J., Houtkamp, R., & Roelfsema, P. R. (2011). Different states in visual working memory: When it guides attention and when it does not. Trends in Cognitive Sciences, 15(7), 327–334. https://doi.org/10.1016/j.tics.2011.05.004

Rose, N. S., LaRocque, J. J., Riggall, A. C., Gosseries, O., Starrett, M. J., Meyering, E. E., & Postle, B. R. (2016). Reactivation of latent working memories with transcranial magnetic stimulation. Science, 354(6316), 1136–1139. https://doi.org/10.1126/science.aah7011

Schneegans, S., & Bays, P. M. (2017). Restoration of fMRI decodability does not imply latent working memory states. Journal of Cognitive Neuroscience, 29(12), 1977–1994. https://doi.org/10.1162/jocn_a_01180

Sprague, T. C., Ester, E. F., & Serences, J. T. (2016). Restoring latent visual working memory representations in human cortex. Neuron, 91(3), 694–707. https://doi.org/10.1016/j.neuron.2016.07.006

Sreenivasan, K. K., Curtis, C. E., & D’Esposito, M. (2014). Revisiting the role of persistent neural activity during working memory. Trends in Cognitive Sciences, 18(2), 82–89. https://doi.org/10.1016/j.tics.2013.12.001

Stokes, M. G. (2015). “Activity-silent” working memory in prefrontal cortex: A dynamic coding framework. Trends in Cognitive Sciences, 19(7), 394–405. https://doi.org/10.1016/j.tics.2015.05.004

van Moorselaar, D., Theeuwes, J., & Olivers, C. N. L. (2014). In competition for the attentional template: Can multiple items within visual working memory guide attention? Journal of Experimental Psychology: Human Perception and Performance, 40(4), 1450–1464. https://doi.org/10.1037/a0036229

Watson, A. B., & Pelli, D. G. (1983). Quest: A Bayesian adaptive psychometric method. Perception & Psychophysics, 33(2), 113–120. https://doi.org/10.3758/BF03202828

Wolff, M. J., Jochim, J., Akyürek, E. G., & Stokes, M. G. (2017). Dynamic hidden states underlying working-memory-guided behavior. Nature Neuroscience, 20(6), 864–871. https://doi.org/10.1038/nn.4546

Xu, Y. (2017). Reevaluating the sensory account of visual working memory storage. Trends in Cognitive Sciences, 21(10), 794–815. https://doi.org/10.1016/j.tics.2017.06.013

Yi, D.-J., Turk-Browne, N. B., Chun, M. M., & Johnson, M. K. (2008). When a Thought Equals a Look: Refreshing Enhances Perceptual Memory. Journal of Cognitive Neuroscience, 20(8), 1371–1380. https://doi.org/10.1162/jocn.2008.20094

Zokaei, N., Heider, M., & Husain, M. (2014). Attention is Required for Maintenance of Feature Binding in Visual Working Memory. Quarterly Journal of Experimental Psychology, 67(6), 1191–1213. https://doi.org/10.1080/17470218.2013.852232

Zokaei, N., Manohar, S., Husain, M., & Feredoes, E. (2014). Causal evidence for a privileged working memory state in early visual cortex. The Journal of Neuroscience, 34(1), 158–162. https://doi.org/10.1523/JNEUROSCI.2899-13.2014

